# Microglia mediate the early-life programming of adult glucose control

**DOI:** 10.1101/2024.07.02.601752

**Authors:** M Valdearcos, ER McGrath, SM Brown Mayfield, A Folick, RT Cheang, L Li, TP Bachor, RN Lippert, AW Xu, SK Koliwad

## Abstract

Mammalian glucose homeostasis is, in part, nutritionally programmed during early neonatal life, a critical window for the formation of synapses between hypothalamic glucoregulatory centers. Although microglia are known to prune synapses throughout the brain, their specific role in refining hypothalamic glucoregulatory circuits remains unknown. Here, we show that microglia in the mediobasal hypothalamus (MBH) of mice actively engage in synaptic pruning during early life. Microglial phagocytic activity is induced following birth, regresses upon weaning from maternal milk, and is exacerbated by feeding dams a high-fat diet while lactating. In particular, we show that microglia refine perineuronal nets (PNNs) within the neonatal MBH. Indeed, transiently depleting microglia before weaning (P6-16), but not afterward (P21-31), remarkably increased PNN abundance in the MBH. Furthermore, mice lacking microglia only from P6-16 had glucose intolerance due to impaired glucose-responsive pancreatic insulin secretion in adulthood, a phenotype not seen if microglial depletion occurred after weaning. Viral retrograde tracing revealed that this impairment is linked to a reduction in the number of neurons in specific hypothalamic glucoregulatory centers that synaptically connect to the pancreatic β-cell compartment. These findings show that microglia facilitate synaptic plasticity in the MBH during early life through a process that includes PNN refinement, to establish hypothalamic circuits that regulate adult glucose homeostasis.

## Introduction

The brain is increasingly recognized as a critical regulator of systemic glucose homeostasis(1), and the intricate circuits involved in this regulation are beginning to be determined. These circuits integrate glucose sensory signaling with multiple effector mechanisms impinging on peripheral tissues controlling blood glucose. For instance, chemogenetically activating AgRP neurons in adult rodents acutely reduces systemic insulin sensitivity(2). Specific glucose-sensing neurons in the hypothalamus also govern the release of insulin and glucagon from the pancreas through the autonomic nervous system(3), and disrupting these mechanisms promotes manifestations of diabetes(4,5). Moreover, recent efforts to understand this brain-to-islet circuitry have identified hypothalamic neurons that synaptically couple to efferent autonomic fibers supplying pancreatic islets(6). This includes a recently uncovered brain-to-β-cell trans-neural circuit regulating insulin secretion involving specific oxytocin neurons in the paraventricular nucleus of the hypothalamus (PVH) that receive input from AgRP neurons comprising the melanocortin system located in the arcuate nucleus (ARC) of the MBH (7).

Circumstances occurring during early neonatal life are important in determining one’s risk of developing diseases such as type 2 diabetes in adulthood (8,9). The hypothalamus is immature at birth and develops postnatally through continuous projection of neuronal axons in the MBH that synapse onto specific targets to establish the melanocortin system. In rodents, for example, AgRP neurons develop axonal projections mainly during the second week of postnatal life (10), sending inputs to the PVH between postnatal day (P) 8 and P10(11). POMC projections display a similar temporal pattern (12). The anatomical and temporal precision of the developmental connections that MBH neurons make with those in other brain regions, are programmed by specific peripheral factors. For example leptin, a hormone the levels of which surge between P8 and P12 in mice, is essential for the developmental wiring of hypothalamic circuits(13); treating leptin-deficient (ob/ob) mice with exogenous leptin until P28 restores neuronal projections that otherwise fail to form, whereas doing so after P28, is relatively ineffective by comparison (14,15).

Nutritional factors also influence CNS control of metabolic function. In particular, obese mothers and those who consume high-calorie diets when their offspring are still neonates increase the risk for an array of metabolic diseases when those offspring reach adulthood(16), a process termed “nutritional metabolic programming.” Similarly, rodents born to obese dams fed a high-fat diet (HFD) during gestation and/or lactation become progressively overweight, hyperphagic, and glucose intolerant as they grow into adults(16,17), metabolic alterations that are associated with disrupted development of AgRP and POMC projections to the PVH. Even when confined to the lactating period, maternal HFD consumption alters the development of hypothalamic neuronal circuits, causing obesity and hyperglycemia in the offspring(18). However, the hypothalamic sensors responsible for detecting hormonal, nutritional, and other metabolic signals during this critical window of susceptibility remain elusive. Similarly, the cellular mechanisms underlying how maternal malnutrition abnormally programs glucoregulatory MBH circuits are poorly understood.

By both genetically deleting key NF-kB regulators and employing chemogenetic strategies specifically in microglia, we recently showed that altering microglial inflammatory responsiveness has a near-immediate effect on systemic glucose homeostasis(19). Specifically, activating microglia improved glucose tolerance in mice fed a HFD, an impact we linked to the regulation of hypothalamic glucose-sensing neurons, a mechanism independent of their previously identified role in regulating energy balance (19). However, the role of microglia in the early-life programming of CNS glucoregulatory circuits has not yet been studied.

Neuronal circuit development entails the formation, maturation, and stabilization of synapses, which consist of presynaptic and postsynaptic elements, the surrounding extracellular matrix (ECM), and glia including microglia. The peptide hormone amylin, which influences neurogenesis, axonal fiber outgrowth, and leptin signaling in the hypothalamus(20–22), also enables the birth of microglia in the ARC(23), suggesting that the emergence of hypothalamic microglia is co-regulated along with hypothalamic circuit development during embryogenesis and early postnatal life. Moreover, depleting fetal microglia, specifically during gestation using the CSF1R inhibitor PLX5622, reduces the number of postnatal POMC neurons in the MBH in association with accelerated weight gain starting at P5(24).

Microglia impact neuronal circuit organization both through synaptic pruning(25–28) and ECM remodeling (29,30). Perineuronal nets (PNNs) are specialized reticular formations within the ECM of the CNS that scaffold neuronal synapses and guide synaptic plasticity, including during development(31). Microglia sculpt PNNs by releasing matrix metalloproteinases and other PNN-degrading proteases(32). Furthermore, recent work in mice shows that microglia remove PNNs in response to neuronally secreted IL-33; blocking this crosstalk impairs microglial phagocytosis and PNN engulfment, leading to impaired spine plasticity, reduced neonatal neuronal input integration, and impaired memory (30). Intriguingly, postnatal PNN formation in the MBH coincides with the critical period for the maturation of AgRP neuronal circuits comprising the melanocortin system(33). Indeed, it was recently shown that depleting microglia during early postnatal life is sufficient to increase PNN density and the overgrowth of AgRP neurons, suggesting that microglia impact the postnatal development of AgRP neurons by controlling PNN formation(34).

In this study, we show that early postnatal microglia help program adult glucose homeostasis in mice by shaping the connectivity between hypothalamic glucoregulatory neurons and the pancreatic β cell compartment, a process that involves an unrecognized microglial phagocytic function guiding postnatal PNN refinement.

## Results

### Microglia exhibit reactivity in the MBH during the early-life period when pups rely on maternal milk

We previously showed that providing milk fat, rich in palmitic acid, but not isocaloric olive oil (oleic acid), to adult mice by enteric gavage is sufficient to induce MBH microglial reactivity. Prompted by this finding, we examined mice during the neonatal period, when they rely on maternal milk similarly rich in palmitic acid for all their nutritional needs. Mice matched for litter size were nursed by dams consuming either a standard low-fat chow diet (LFD) or a HFD and underwent hypothalamic dissection at different time points before weaning (P8 and P16, respectively) or one week after weaning (P28). Remarkably, we observed that both the number of microglia and the size of their cell bodies increased markedly from P8 to P16 specifically in the MBH (Figures 1A-C), but not in in adjacent hypothalamic areas such as the VMH, lateral hypothalamic area (LHA), dorsomedial hypothalamus (DMH), or in other brain regions such as the cerebral cortex (Figure S1). Moreover, the response was transient, with a substantial regression seen by P28 after the mice began foraging for chow instead of consuming maternal milk (Figures 1A-C). Additionally, this transient MBH microglial response was accentuated when the dams consumed a HFD while lactating (Figures 1D-F). Together, these findings prompted us to examine the cellular processes that might enable these microglia to influence MBH circuit development.

**Figure 1.**
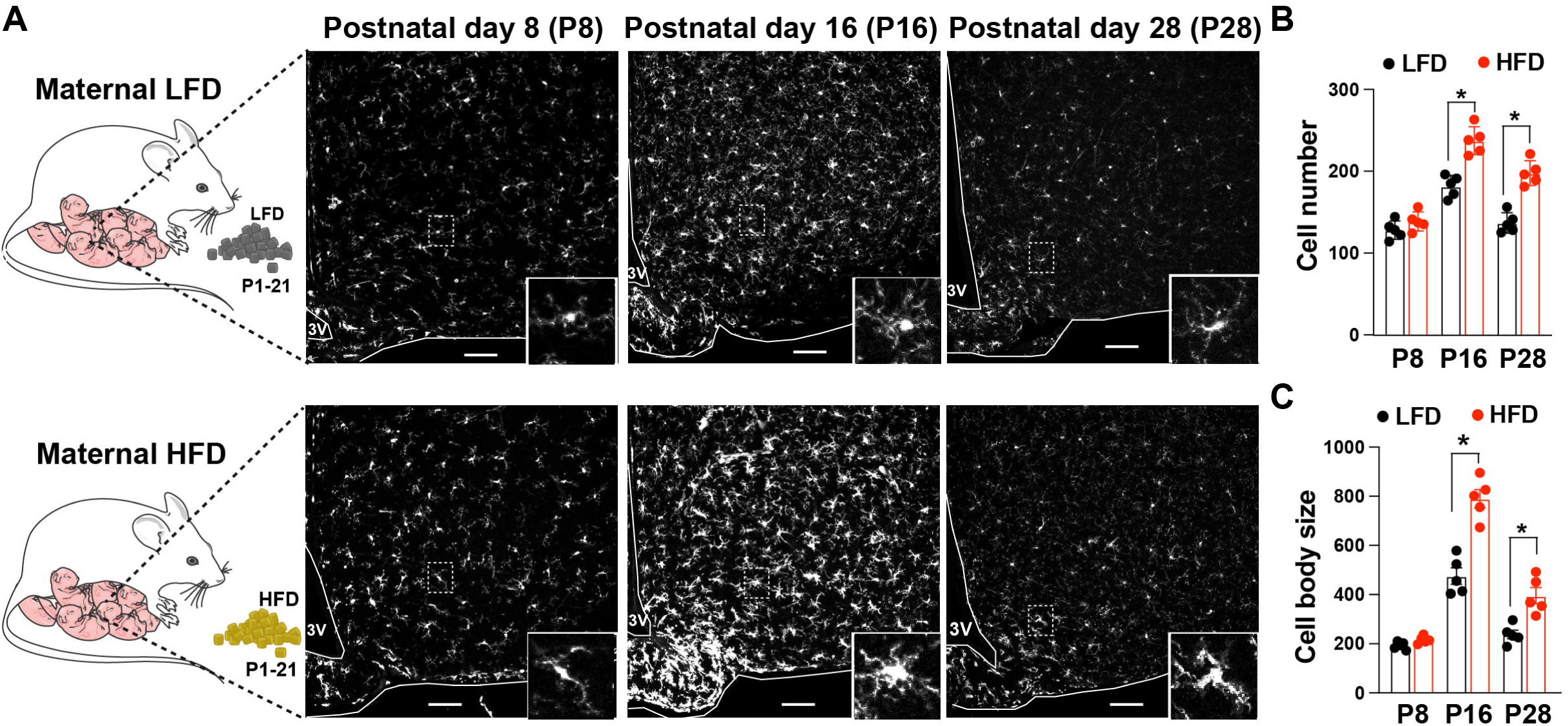
Maternal high-fat diet during lactation alters the offspring’s hypothalamic microglial morphology and reactivity status. A) Representative confocal images postnatal MBH sections, showing the increased number and cell body size of Iba1+ microglia at postnatal day 16 with dams fed a standard low-fat diet (LFD) or high-fat diet (HFD) during lactation. These changes are sustained after weaning at postnatal day 28 when mice are fed a low-fat diet since postnatal day 21. B) Quantification of cell number in A. C) Quantification of cell body size in A (n= 5-6/group; * P< 0.05 HFD vs LFD). Values are mean ± SEM (3V, third ventricle: scale bar, 20 μm).

### Maternal HFD affects microglial synaptic engulfment and the formation of PNNs during the postnatal development of hypothalamic neuronal circuits

Microglia are crucial for postnatal synaptogenesis, working to eliminate weak or unnecessary synapses by selectively engulfing synaptic material for degradation in the lysosomal compartment(35,36). Given that the synaptic connections forming the hypothalamic circuits controlling peripheral metabolism coincide, both spatially and temporally, with the microglial reactivity we observed in the MBH of mice prior to weaning, we examined the involvement of these microglia in synapse elimination. To do so, we used 3D Imaris reconstructions of MBH microglia to visualize internalized synaptic material marked by lysosomal postsynaptic density protein 95 (PSD-95) (Figures 2A-B). Interestingly, both overall microglial phagocytic activity and PSD-95 engulfment in the MBH peaked at P16 and decreased after weaning (P28), indicating that these reactive microglia were engaged in synaptic refinement. Notably, these indicators of microglial phagocytosis and synaptic engulfment were further accentuated at P16, and the post-weaning regression at P28 was less profound, when the dams were fed a HFD (Figures 2C-D), suggesting that maternal HFD consumption both heightens and prolongs the demand on microglia to refine the synapses of MBH circuits governing metabolic homeostasis. Taken together, these findings highlight the importance of microglia in mediating the interplay between maternal nutrition and synaptic plasticity within the MBH during early life.

**Figure 2.**
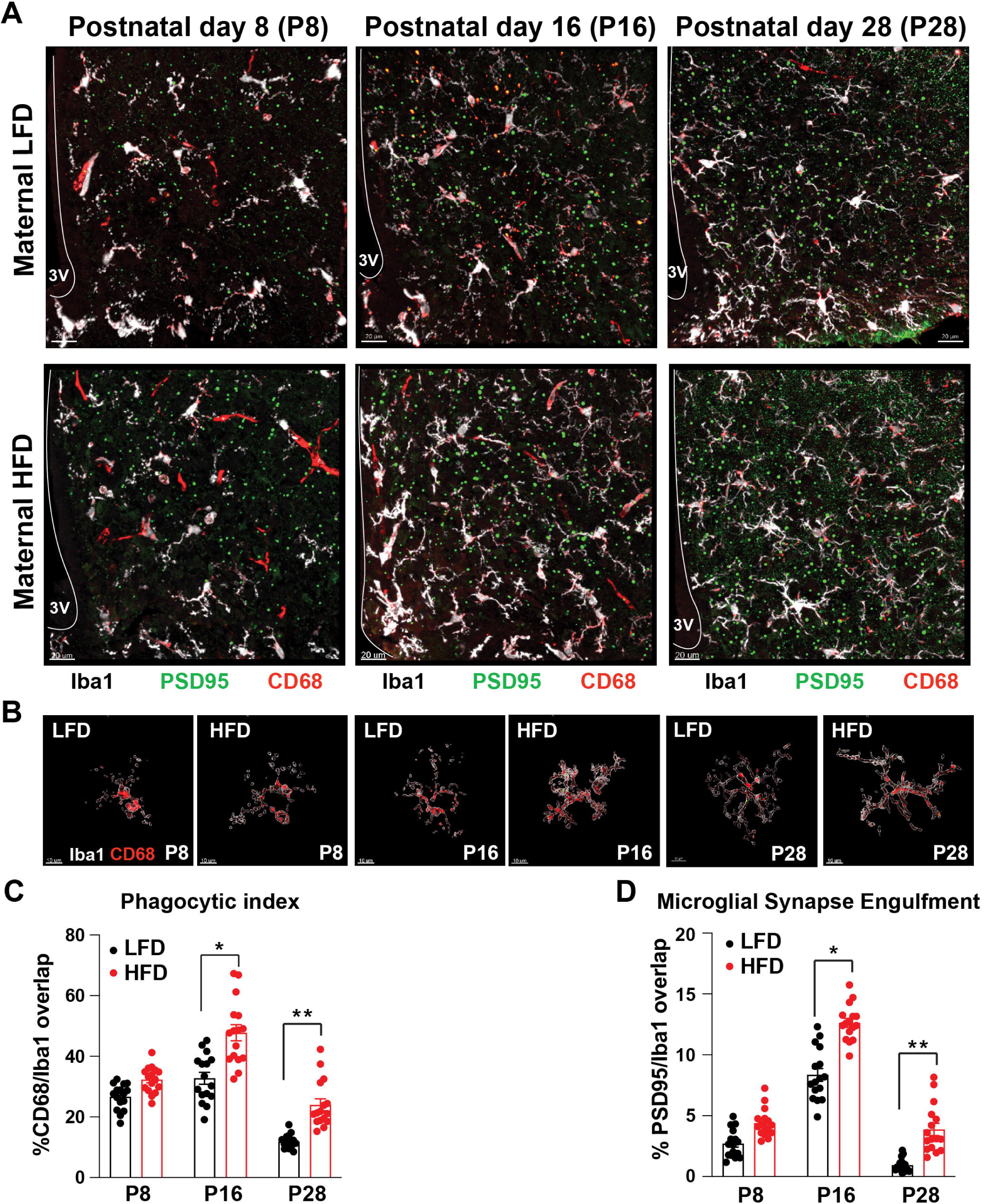
Maternal high-fat diet affects microglial synaptic engulfment during the postnatal development of hypothalamic neuronal circuits. A) Representative confocal images of postnatal MBH sections, showing Iba1+ microglia (white), PSD-95 (green, postsynaptic density protein 95), and CD68 (red, lysosomal content) with dams fed a standard low-fat diet (LFD) or high-fat diet (HFD) during the lactation period. B) Representative 3D surface Imaris reconstruction of microglia with lysosomal content (CD68, red) in microglial cells at P8, P16, and P28. C) Quantification of CD68 content in microglial cells. E) Quantification of microglial engulfment of PSD-96 (n=3 mice/condition, n=6 cells/section, a total of 48 cells analyzed. Values are mean ± SEM (3V, third ventricle: scale bar, 20 μm).

Given that the maturation of AgRP neurons during the lactation period coincides with the formation of PNNs(33), we also investigated the extent to which microglia remodel PNNs during this period. Consistent with previous studies(33), we found that PNNs labeled with wisteria floribunda agglutinin (WFA) began appearing in the ARC between P8 and P16 and increased further by P28 (Figure 3A). Moreover, microglia in the ARC during this time were seen to engulf aggrecan, a core PNN protein, within CD68^+^ lysosomes (Figure 3B). As was observed for synaptic engulfment, PNN engulfment by MBH microglia during the neonatal period was increased when the dams were fed a HFD (Figure 3C). These data, together with our earlier findings, indicate that maternal diet dually influences microglial synaptic and PNN engulfment within the MBH during a critical window when the hypothalamic circuits that regulate peripheral metabolic homeostasis are developing.

**Figure 3.**
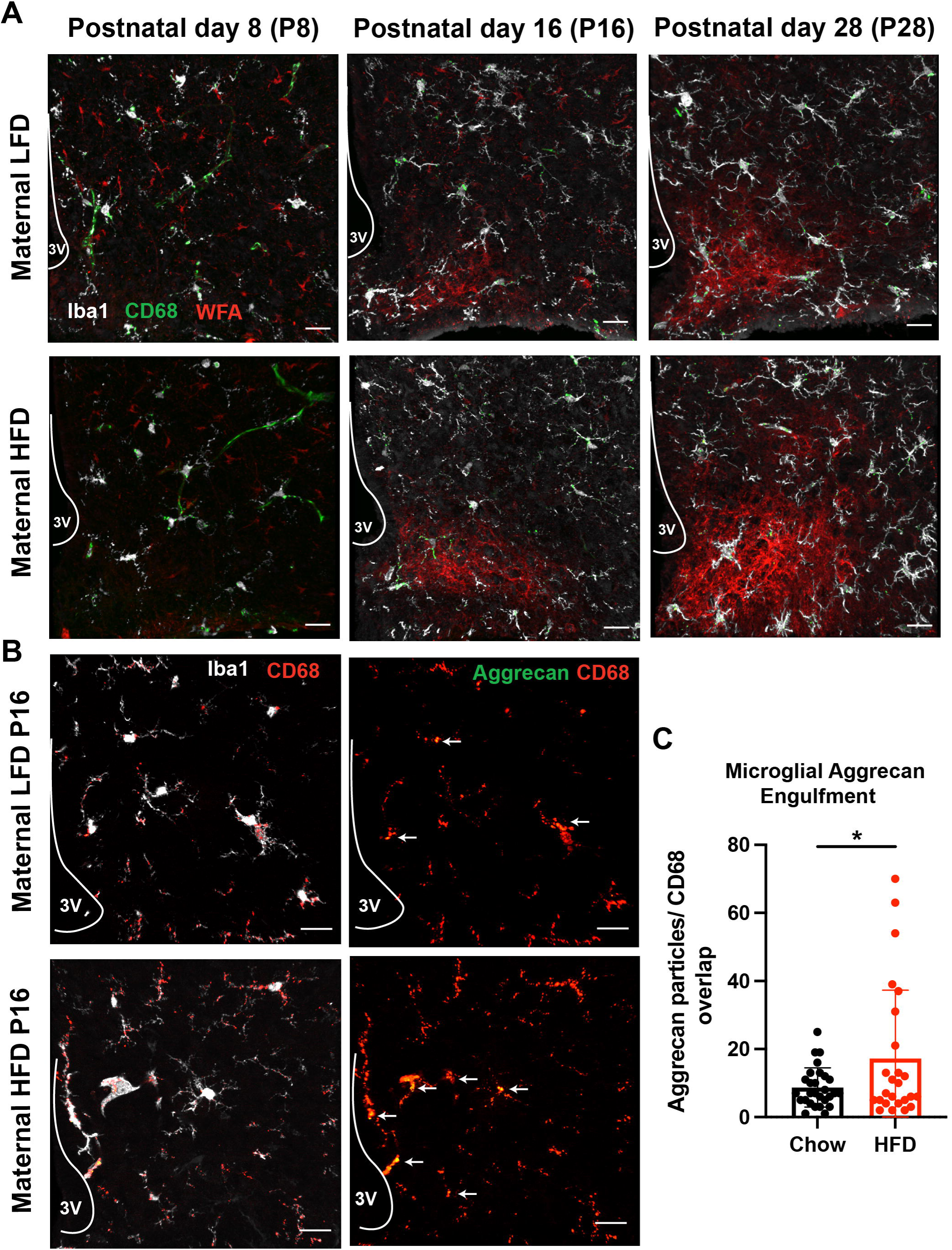
Maternal high-fat diet increases microglial aggrecan engulfment during the postnatal formation of perineuronal nets. A) Representative confocal images of postnatal MBH sections, showing Iba1+ microglia (white), CD68 (green, lysosomal content), and WFA (red, Wisteria floribunda agglutinin to label perineuronal nets) with dams fed a standard low-fat diet (LFD) or high-fat diet (HFD) during the lactation period. B) Representative confocal images of postnatal MBH sections at postnatal day 16, showing Iba1+ microglia (white), Aggrecan (green, chondroitin sulfate proteoglycan of perineuronal nets). C) Quantification of microglial engulfment of Aggrecan in lysosomal content (n=5 mice, n=5-6 sections/mice, each dot represents the overlap of aggrecan particles with CD68 per section). Values are mean ± SEM (3V, third ventricle: scale bar, 20 μm).

### Depleting microglia specifically during early life enforces a build-up of PNNs in the ARC and promotes glucose intolerance in adulthood

Based on these findings, we hypothesized that microglia may determine PNN density within the MBH during early postnatal life. To explore this, we used an inhibitor of CSF1 receptor signaling (BLZ945, 200mg/kg, ApexBio) to effectively and reversibly deplete CNS microglia in mice specifically before weaning from P6 to P16, with microglia fully repopulating the brains of mice within 5 days of discontinuing BLZ945 treatment (Figures 4A-B). Doing so clearly increased the density of PNNs within the MBH, underscoring the importance of microglia in sculpting these PNNs during early-life hypothalamic circuit development(Figures 4C-D).

**Figure 4.**
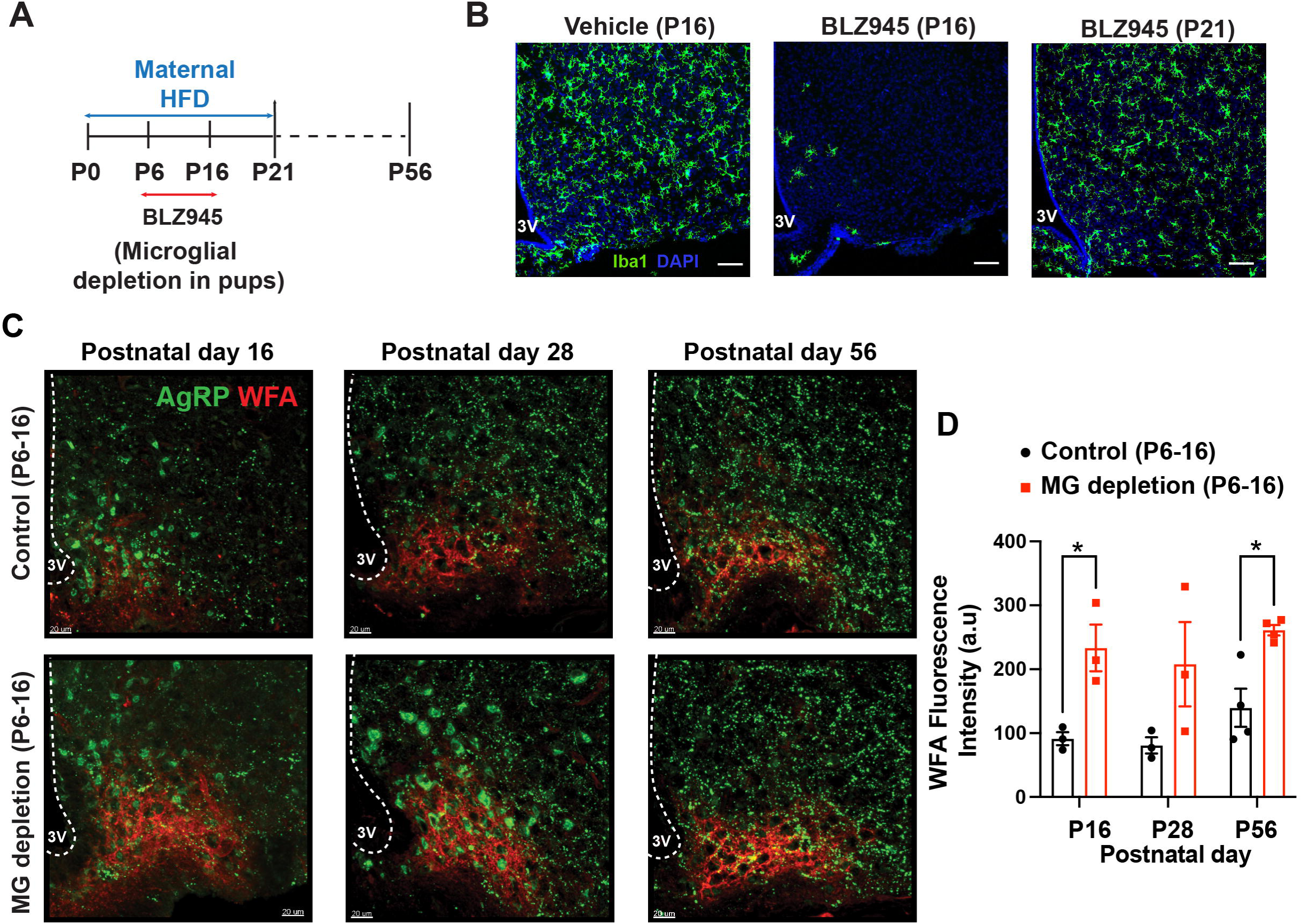
Transient postnatal microglial depletion enhances hypothalamic perineuronal net abundance in the offspring of HFD-fed dams. A) Experimental design to assess the impact of postnatal microglial depletion from P6-16 using BLZ945 treatment (200mg/kg, every other day, i.p.) while dams are fed a high-fat diet during lactation. B) Representative confocal images of MBH sections, showing Iba1+ (green) microglia repopulation after drug withdrawal (P16). C) Representative confocal images showing cell bodies and axonal projections of AgRP (green) neurons and WFA (red, Wisteria floribunda agglutinin to label perineuronal nets) at P16, P28, and P56. D) Quantification of WFA fluorescence intensity in C (n=3 mice/condition). Values are mean ± SEM (3V, third ventricle: scale bar, 20 μm).

We further hypothesized that the overabundance of PNNs in the MBH at the conclusion of the neonatal period stemming from this transient impairment of microglial sculpting might lead to abnormal programming of CNS control over peripheral metabolism. In testing this hypothesis, we noted that as in our prior studies involving adult mice fed a HFD(37), microglial depletion prior to weaning was associated with the pups gaining less weight while their microglia were depleted vs. control mice (Figure 5A). Interestingly, upon discontinuing BLZ495 treatment and in association with the reconstitution of the microglial compartment, the weights of the young mice rapidly normalized to those of control mice that had retained their microglia throughout the study. However, despite the fact that both groups of mice had similar weight trajectories thereafter and had similar weights when analyzed in adulthood (8 weeks of life), the mice that underwent microglial depletion from P6-16 were remarkably glucose intolerant as adults vs their control counterparts (Figures 5B-C). These data indicate that neonatal microglia are required to ensure normal systemic glucose tolerance in adulthood, an outcome that is not a function of differences in body weight gain.

**Figure 5.**
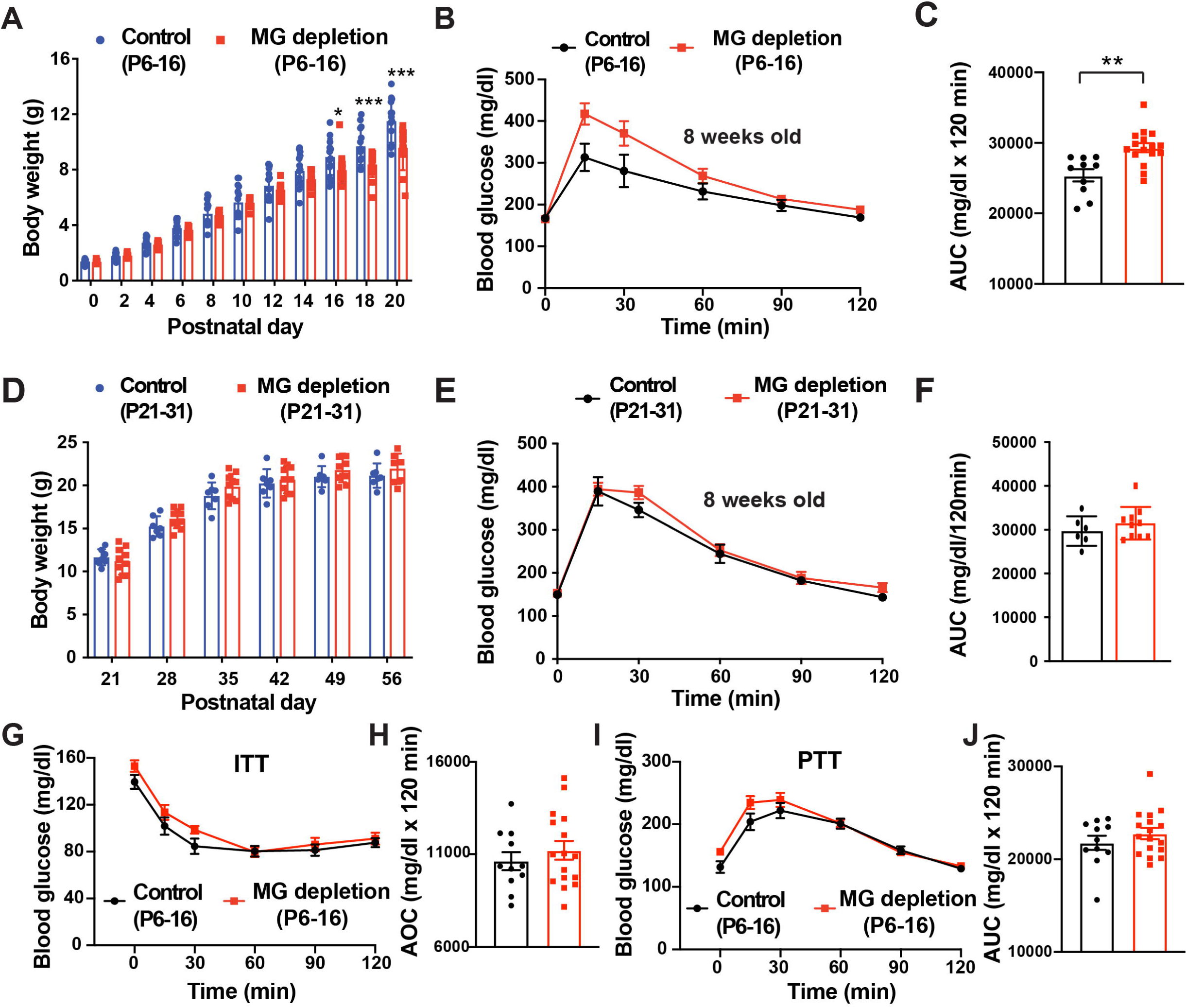
Depleting microglia specifically during the neonatal period (P6-16) induces glucose intolerance in adult offspring of HFD-fed dams. A) Reduced body weight and B) worsened glucose tolerance test (GTT) at 8 weeks of age in microglial-depleted pups from P6-16 when dams were fed a high-fat diet during lactation. C) Area under the curve (AUC) of B (n=10-16/group). D) Equivalent body weight and E) glucose tolerance when microglia are depleted from P21-31 when dams were fed a high-fat diet during lactation. F) AUC of E (n=6-10/group). G) Equivalent insulin tolerance test at 9 weeks of age in microglial-depleted pups from P6-16 when dams were fed a high-fat diet during lactation. H) Area over the curve (AOC) of G (n=11-16/group). I) Equivalent pyruvate tolerance test (PTT) at 10 weeks of age in microglial-depleted pups from P6-16 when dams were fed a high-fat diet during lactation. J) AUC of I (n=11-16). Values are mean ± SEM.

To confirm the temporal importance of microglia in this aspect of metabolic programming, we examined a separate cohort of mice in which microglia were similarly depleted for a 10-day window, but after weaning had already occurred (P21-31). Doing so showed that unlike the clear impact of depleting microglia from P6-16, depleting microglia from P21-31 affected neither weight gain during the period of the depletion nor the glucose tolerance of the mice when they were assessed at eight weeks of life (Figure S3). This lack of effectiveness was not a function of differences in the potency of CSF1 inhibition based on the age of the mice. Indeed, CSF1 inhibition depleted microglia in the brains of mice just as effectively from P21-31 as from P6-16 (Figure S3B). These data also strongly suggest that our findings were not due to aberrant populations of microglia emerging in the brain following drug-induced depletion, as microglial repopulation occurred similarly following depletion both from P6-16 and P21-31 but only depletion from P6-16 impacted the metabolic parameters we analyzed. To further convince ourselves that our findings were on-target, we also treated mice from P6-16 with PLX73086, a different CSF1R inhibitor that does not penetrate the CNS(38). Such treatment neither depleted microglia nor affected adult glucose tolerance (Figure S4), indicating that our prior results were not a function of depleting non-CNS myeloid populations. Taken together, our findings indicate that microglia are required during early neonatal life to ensure normal glucose tolerance in adulthood, highlighting the possibility that microglia may play a role during early life in programming the capacity of the CNS to regulate glucose homeostasis later in life.

Given that low birth weight, including in humans, followed by catch-up growth during infancy, can negatively impact glucose homeostasis and the risk for type 2 diabetes in adulthood (39,40), we also sought to ensure that our findings were not a function of the rapid weight normalization experienced by mice following neonatal microglial depletion. In examining this, we noted that neonatal microglial depletion impacted male and female mice differently. In both male and female mice, microglial depletion from P6-16 led to a slight reduction in weight gain during the period of depletion, followed by rapid body weight normalization upon discontinuation of CSF1R inhibition (Figure S5). By contrast, whereas depleting microglia from P6-16 caused adult glucose intolerance in male mice, this was not clearly seen in female mice (Figure S5). This sex-dependent difference, a topic for future studies, indicates that our findings in male mice are not simply a consequence of reduced neonatal weight gain followed by catch-up growth, as this phenomenon occurred in female mice as well. Additionally, although early-life microglial depletion impacted neonatal weight gain, it did not affect linear growth. This detail is important, as small-for-gestational-age babies who later exhibit impaired glucose metabolism usually have both reduced weight and length at birth.

Intriguingly, the impact of eliminating neonatal microglia on adult glucose intolerance phenotype was clearly evident when evaluating mice that had been nursed by dams consuming a HFD while lactating, but not as obvious when examining mice that had been nursed by dams fed a fat-sparing LFD (Figure S2). This finding is concordant with our assessments of microglial number and phagocytic activity within the MBH, which were both much higher during neonatal life in the context of maternal HFD, and suggests that maternally derived nutritional components, potentially including fats, may be important in driving microglia to engage in the developmental programming of metabolic circuits within the hypothalamus.

### Microglia act during early life to ensure normal adult glucose-responsive pancreatic insulin secretion

*Next, we sought to identify the determinant of systemic glucose homeostasis that is programmed by early-life microglia*. First, we noted that although depleting microglia (P6-16) worsened glucose tolerance, it did not affect insulin tolerance (ITT) (Figures 5 G-H) or pyruvate tolerance (PTT) (Figures 5I-J), suggesting that our findings were not due to either impaired systemic insulin sensitivity or hepatic glucose production. By contrast, mice lacking microglia from P6-16 had lower plasma insulin levels throughout GTTs performed at 8 weeks of life than did controls, suggesting the possibility of reduced glucose-responsive pancreatic insulin secretion (Figures 6 A-B). We explored this possibility by examining both the β-cell mass and glucose responsiveness of pancreatic islets from pancreata isolated at 8 weeks of life from control mice vs. those having undergone microglial depletion from P6-16. We found that despite a clear loss of systemic glucose tolerance, the pancreatic islets of adult mice having undergone microglial depletion during early neonatal life had both a similar β-cell mass within the pancreas (Figure 6 C) and displayed similar glucose-stimulated insulin secretion (GSIS) dynamics (Figures 6 D-H) as did those of control mice. Together, these findings suggest that if early-life microglia programs the ability of the CNS to regulate adult pancreatic glucose responsiveness, it does not do so by impacting either β-cell development or functional maturation *per se*. This conclusion and work indicating that autonomic control of glucose homeostasis is programmed during early life by maternal nutritional factors(41,42), led us to explore instead the likelihood that microglia program the autonomic connectivity between the hypothalamus and the pancreatic ý-cell compartment.

**Figure 6.**
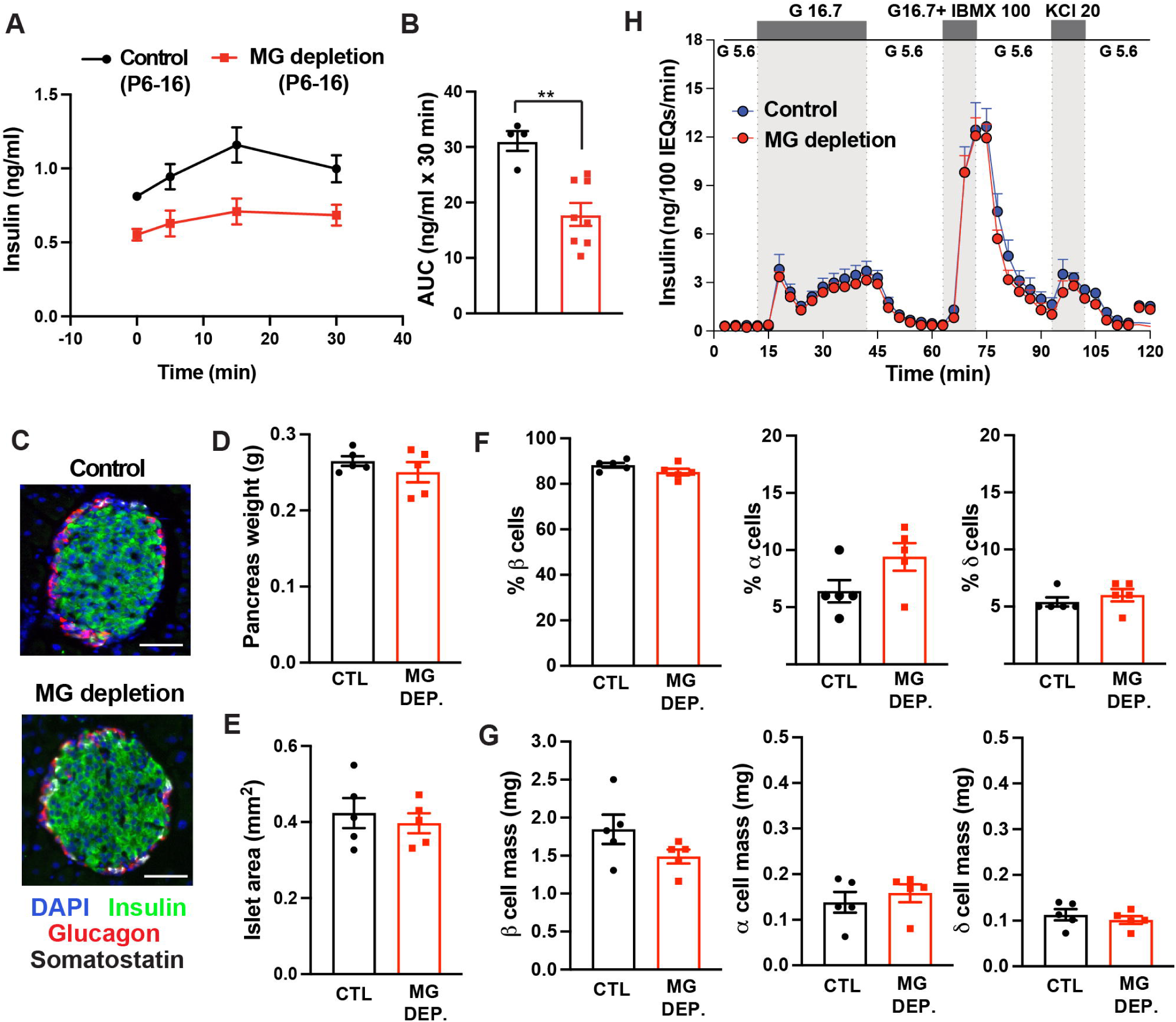
Microglial depletion during the neonatal period (P6-16) results in defective insulin secretion without affecting the development and cell-intrinsic features of β cells. A) Reduced insulin secretion in vivo in microglial-depleted mice (P6-16) after i.p glucose injection (2g/kg) in 8-week-old mice. B) Area under the curve (AUC) of A (n=4-8/group). C) Representative images of pancreatic islets showing DAPI (blue), insulin (green), glucagon (red), and somatostatin (white) in microglial-depleted mice (P6-16) and control mice from 8-week-old mice. D) Equivalent pancreas weight, E) islet area, F) cellular composition, and G) cellular mass and H) glucose-stimulated insulin secretion by isolated islets in microglial-depleted mice (P6-16) compared with control mice at 8 weeks of age (n=5/ group, ** P< 0.01). Values are mean ± SEM, scale bar, 50 μm.

### Early-life microglia program the autonomic connectivity between the β-cell compartment and the hypothalamus, including known glucoregulatory neuronal populations

With this question in mind, we developed a strategy to fluorescently trace the synaptic connectivity between the pancreatic β-cell compartment and hypothalamic neurons, which are joined by autonomic fibers that communicate with key hypothalamic glucoregulatory centers(6,7). We used a new-generation pseudorabies virus (PRV-Ba2001), which expresses a Cre-inducible GFP reporter and the thymidine kinase enzyme, injected (10^8^pfu/ml) via a minimally invasive surgical approach into the pancreatic head, body, and tail regions of MIP-Cre^ERT^ mice which express Cre specifically in β-cells (Figures 7A-B) as described previously (21). Upon subsequent tamoxifen treatment, this model enabled us to trace GFP fluorescence from the β-cell compartment through neurons, including those comprising the autonomic nervous system, across multiple ascending synapses into specific hypothalamic nuclei including the ARC and PVH. Using this approach, we observed that transiently eliminating microglia in neonatal mice from P6-16 was remarkably sufficient to reduce the number of neurons within the ARC and the PVH that were specifically connected polysynaptically to the β-cell compartment (GFP^+^) at eight weeks of age (Figures 7C-G). In exploring the identity of these β-cell-connected hypothalamic neuronal populations, we found that those numerically reduced in adulthood by neonatal microglial depletion remarkably included AgRP neurons in the ARC (Figure 7D), which are known to be crucial for CNS glucose regulation and are already implicated in the impairment of glucose homeostasis induced by maternal HFD consumption during early life(18). These data indicate that microglia during the early neonatal period help program the ability of the adult CNS to properly potentiate glucose-responsive pancreatic insulin secretion by determine the connectivity of hypothalamic neurons, including those involved in glucose sensing, with the pancreatic β-cell compartment.

**Figure 7.**
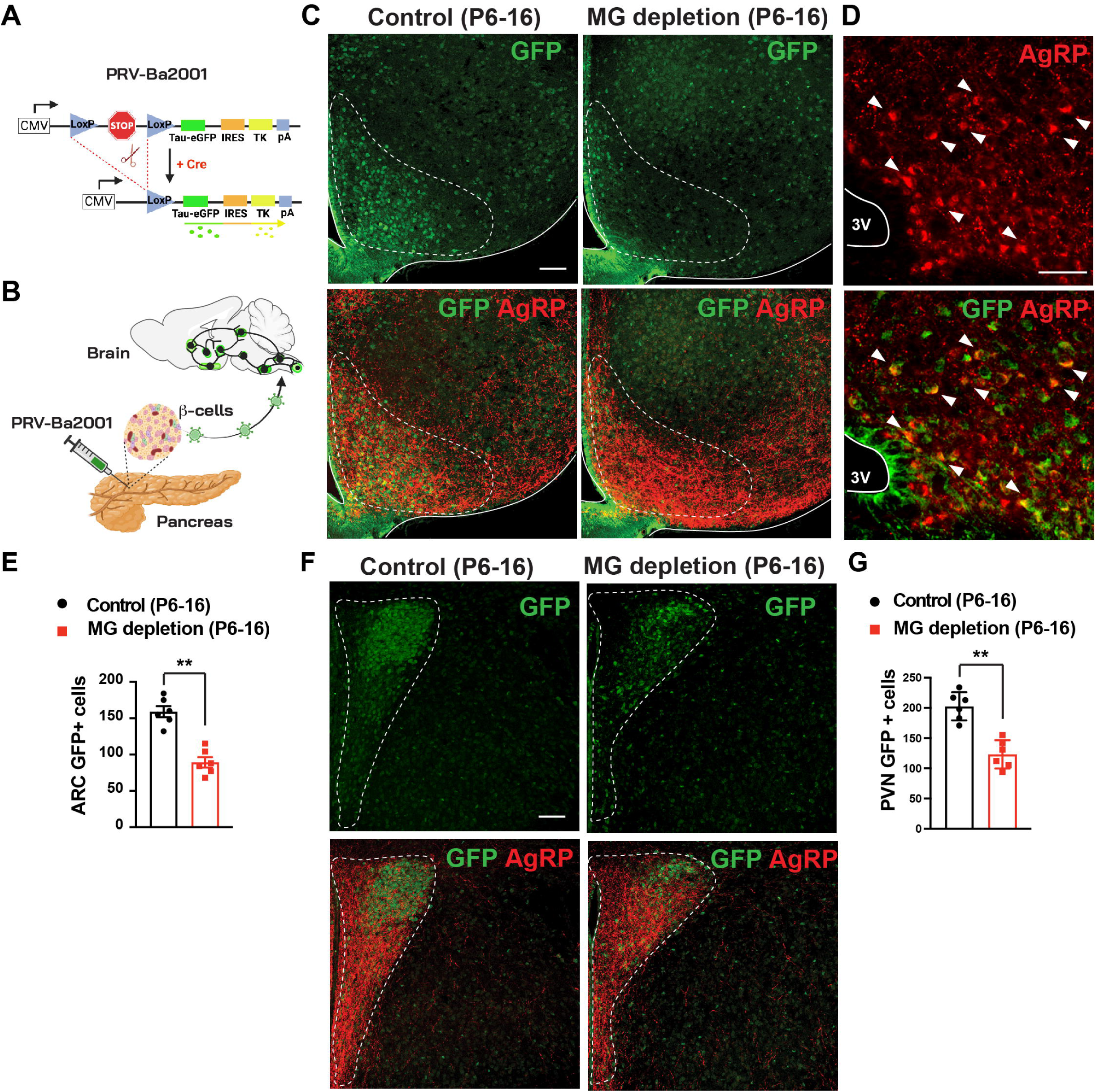
Microglial depletion during the neonatal period (P6-16) impairs the synaptic connectivity between hypothalamic neurons and β cells. A) Structure and expression properties of pseudorabies virus (PRV) strain Ba2001 that is dependent on Cre-mediated recombination for the expression of GFP and thymidine kinase (TK), a viral protein that promotes efficient replication. B) Schematic representation of Ba2001 delivery into the pancreas of MIP-CreER mice to target insulin-expressing β cells and retrogradely trace efferent neurons from the brain using GFP expression. C) Representative images of MBH sections showing GFP + neuronal cells (green) synaptically connected to β cells in the arcuate nucleus. D) Co-localization between GFP and AgRP cell bodies (red). E) Quantification of D (n=6/group,** P< 0.01). F) Representative images showing GFP + neuronal cells (green) synaptically connected to β cells in the paraventricular nucleus of the hypothalamus (PVH). G) Quantification of F (n=6/group,** P< 0.01). Values are mean ± SEM (3V, third ventricle: scale bar, 20 μm).

## Discussion

In this study we show that in mice, microglia are reactive within the MBH during early neonatal life, working to prune synapses and sculpt PNNs within this region of the brain. Building on this, we reveal a previously unrecognized role for microglia during the early neonatal period, when mammals rely on maternal milk, in programming the synaptic connectivity of hypothalamic neurons with the pancreatic β-cell compartment. Moreover, our work implicates this process as a determinant of the normal adult capacity to elevate insulin levels in response to rising systemic glucose concentrations.

Hypothalamic microglial reactivity was initially described in obese mice and humans over a decade ago(43,44). At the time, such reactivity was interpreted as a detrimental response to diet-induced hypothalamic injury, likely stemming from the classical observation that innate immune cells accumulate at sites of injury or infection and the finding that lipid accumulation, including long-chain saturated fats, can drive a pro-inflammatory reaction in tissue myeloid cells(45,46). Our understanding has significantly evolved from these early studies, with an increasing recognition of the ability of microglia to influence hypothalamic function in both physiologic and pathologic states. For instance, we recently demonstrated that inducing microglial inflammatory signaling within the hypothalamus of both lean and obese adult mice rapidly and unexpectedly benefits glycemic control by enhancing glucose disposal through a melanocortin-dependent pathway regulating pancreatic insulin secretion(19). The present study indicates that microglia are not only involved in acutely modulating CNS control over systemic glucose homeostasis during adult life, but also help to shape the synaptic connections that form during early life in order to enable the capacity for such efferent control well after weaning. In this way, our work bolsters the emerging notion that microglia are assistive, rather than injurious, cells within the context of the developing hypothalamus, a concept that has implications not only for the early life period, but for our interpretation of the role of microglia within the adult hypothalamus as well.

Microglia are increasingly being recognized as critical regulators of neuronal numbers and the refinement of neural circuits during brain development through multiple mechanisms such as releasing factors with neurotrophic functions to support neuronal survival and contributing to the establishment of neuronal wiring in the postnatal brain(47,48). In addition, microglia contribute to the maturation and refinement of synaptic networks in the postnatal period. Our data indicate that postnatal microglia around P16 exhibit enlarged cell bodies and heightened phagocytic activity with increased PSD-95-positive puncta engulfment in the MBH, coinciding with the development of AgRP and POMC axonal projections(11). Previous studies have shown that “ eat me signals,” such as complement cascade components C1q and C3, are essential components to promote microglial engulfment of inappropriate and/or extraneous synapses through the microglial C3 receptor(49,50). In contrast, “don’t eat me” signals such as CD47 serve as a negative regulator of synaptic pruning to avoid excessive elimination of synapses(51). Further studies are needed to determine the signals by which maternal diet influences microglial phagocytic activity during the development of hypothalamic neural circuits.

Microglia also influence synaptic connectivity by modifying the extent to which the ECM allows neuronal projections to reach their desired targets. PNNs are unique reticular ECM formations that encapsulate the soma and dendrites of certain neurons in the brain, such as fast-spiking parvalbumin-positive inhibitory neurons. PNN formation occurs during critical periods of developmental plasticity, and microglia are integral in remodeling PNNs in both healthy and diseased states(29), regulating the balance between their formation and degradation via both direct and indirect enzymatic/phagocytic processes(52–54). Notably, PNN deficiency in the ARC of postnatal ob/ob mice can be restored by leptin administration during postnatal development(33), and recent work showed that chemically disrupting PNNs in the MBH attenuates the glucose-lowering effects of centrally acting fibroblast growth factor 1(FGF 1) in obese Zucker diabetic fatty (ZDF) rats(55), indicating a potential role of PNNs in regulating glucose homeostasis. Here, we show that postnatal microglial depletion enforces an accumulation of PNNs in the MBH, findings that align with a recent study suggesting that microglia influence AgRP postnatal development by remodeling PNN-dependent synaptic plasticity(34). This is important, given that PNNs selectively enmesh AgRP neurons in the MBH, and their formation coincides with the critical period of AgRP neuron maturation during postnatal development that is influenced by leptin (33).

Synaptic tracing studies and chemogenetic manipulations targeting distinct subpopulations of hypothalamic neurons have provided new insights into how the brain regulates the β-cell insulin secretory response to hypoglycemia(7). For example, a subpopulation of oxytocin neurons in the PVN was recently revealed to comprise a circuit activated by glucoprivation that suppresses insulin secretion by pancreatic β cells via the sympathetic autonomic nervous system. Another recent study by Bruning and colleages used optogenetic and chemogenetic approaches to show that AgRP neuronal activation induces hepatic autophagy as part of a hypothalamic-liver axis that is metabolically adaptative during nutrient deprivation(56). Here, we map brain-to-β-cell synaptic connections as a means to reveal that early postnatal microglia are required for the functional development of glucoregulatory circuits emanating from the MBH that are connected to the β cell compartment. Although our analysis identifies projections from AgRP neurons in both the ARC and PVN as being among those shaped by microglia during early life, future studies will need to focus on additional neuronal subsets across a broader array of brain regions. In addition, we do not know the extent to which microglia are also required for the programming synaptic connectivity between the brain and other peripheral metabolic organs, such as liver, that regulate glucose homeostasis.

The alterations in CNS programming that lead to increases in the risk for cardiometabolic disease later in life displays sex-dependent skewing(57). For example, data from mice suggest that female pups are more susceptible to developing impaired glucose tolerance as adults in response to programming during gestation(58). However, the mechanisms underlying these and other sex-based differences are not understood. In our study, both male and female mice displayed a similar modest body weight reduction during microglial depletion, followed by rapid catch-up growth after drug withdrawal and consequent microglial repopulation of the brain. However such transient microglial depletion, when performed prior to weaning, impacted adult glucose homeostasis only in the male mice we examined despite the fact that our strategy to deplete microglia was equally effective in both sexes. With this in mind, we note that there is a growing appreciation of sex differences in microglial differentiation within the brain(59). Indeed, there are sex differences in microglial gene expression and phagocytic activity during brain development (60,61). Additional studies are needed to understand the sex-specific ways by which microglia shape glucoregulatory synapses within the hypothalamus during early life.

In summary, we show that prior to weaning, microglia shape PNNs within the MBH and that depleting microglia, particularly in the context of a maternal HFD, notably increases PNN density during this critical early-life period. PNN overabundance in the MBH due to microglial depletion, in turn, reduces the connectivity between key hypothalamic glucoregulatory neurons and pancreatic ý cells, the consequence of which is impaired glucose-responsive insulin secretion and relative glucose intolerance in adulthood.

## Limitations of the study

We acknowledge some limitations of this study. First, we do not know if the microglia repopulating the brain after depletion are functionally similar to the microglia in the brains of untreated control mice. However, transcriptional studies suggest that brain homeostasis is not significantly influenced by post-ablative microglial repopulation, suggesting that original resident and repopulated microglia function similarly(62). This concept is underscored by the fact that microglial depletion from P21-31 in our study did not impact adult glucose homeostasis even though such depletion was also associated with repopulation. Second, our studies alone do not prove that the reduced brain-to-β-cell synaptic connectivity in adult mice following neonatal microglial depletion is the specific cause of the impaired adult glucose tolerance we observed. Indeed, this is exceptionally difficult to determine without a temporal gain-of-function approach to independently restore synaptic connectivity despite early life microglial depletion. Current tools and techniques preclude such a definitive study, although further studies are warranted to profile a wider array of potential brain glucoregulatory circuits regulated by microglia during neonatal period. Additionally, our studies do not prove that diminished brain-to-β-cell synaptic connectivity is a direct consequence of altered PNN density within the MBH. Although studies have effectively used enzymatic approaches to diminish PNNs in anatomically defined regions of the brain in adult mice(55), such approaches are not feasible in early neonatal mice. Still, the literature has strongly implicated PNNs in modulating hypothalamic neuronal projections, including those regulating blood glucose(55), and so our conclusion is, in our view, logical. Finally, we do not know that the reduction in neuronal connectivity between the MBH and β-cell compartment induced by early-life microglial depletion is specifically responsible for reduced insulin levels and impaired glucose tolerance in adult mice. Proving this would require independently reestablishing this connectivity in the face of microglial depletion, something that is not yet scientifically approachable.

## Experimental procedures

### Mice

All studies used C57BL/6 background mice. For PRV-Ba2001 tracing, MIP-CreERT(JAX, 024709) adult mice (10-14 weeks of age) were used, where nuclear Cre expression is restricted to pancreatic b-cells when induced by tamoxifen treatment as previously described(6). The appropriate progeny resulting from crosses of these mice, and corresponding controls, received three 5 mg doses of tamoxifen (MP Biomedicals, 10540-29-1) dissolved in 200 μl warm purified corn oil by enteric gavage in order to induce Cre expression. All mice were group-housed and age-matched with *ad libitum* access to water and a specified diet in a pathogen- and temperature-controlled room with a 12:12h light: dark cycle. Mice were fed either a standard low-fat CD (LabDiet 5053) or a HFD (42% fat (kcal), TD. 88137; Envigo. Microglia were depleted in adult mice by administering the CSF1R inhibitor PLX5622 (Plexxikon), formulated in either a low-fat CD (AIN-76A, Research Diets) at a dose of 1.2 g/kg. Neonatal mice received subcutaneous injections of BLZ945 (APExBIO, B4899, 200 mg/kg, every other day), a small molecule inhibitor of CSF1R, dissolved in 20% captisol (Cydex, RC0C7100, beta-cyclodextrin sulfobutyl ethers, sodium salts) to eliminate microglia as described previously (63). Anesthesia was by isoflurane, 100mg/kg ketamine and 10 mg/kg xylazine, or Avertin (terminal procedures). All procedures were performed in accordance with NIH Guidelines for Care and Use of Animals and were approved by the Institutional Animal Care and Use Committees at the University of California San Francisco.

### PRV Neuronal tracing

PRV-Ba2001 was acquired by PNI (Princeton Neuroscience Institute) Viral Vector Core facility. A total of 10ul of PRV-Ba2001 vector (10^8^ plaque-forming units [pfu]/ml) was injected in three separate sites in the head, body and tail regions of a mouse pancreas by intra-abdominal surgery as described previously(6). Animals were sacrificed 120h postsurgery and brains were harvested, fixed in 4% (v/v) paraformaldehyde, and embedded in M-1 Embedding Matrix (Epredia^TM^ 1310).

### In vivo Assesment of Glucose Homeostasis

Body composition analysis including fat mass and lean mass measurements was performed at the NIDDK-funded Nutrition Obesity Research Center (NORC) metabolic core at UCSF, using EchoMRI in awake, concious mice. Glucose tolerance tests (GTT) were performed as described previously. Briefly, animals were fasted for 6h (09:00-15:00) prior to glucose administration (D-(+)-glucose, 20% solution, 2g/kg, i.p: Teknova). Glucose was measured in tail blood by hand-held glucometer. Total glucose area-under-curve (AUC) and excursion was measured by the trapezoid rule. Glucose-stimulated insulin secretion was performed as in the GTT with minor modifications. Glucose (2mg/kg/lean mass i.p.) was administered with collection of tail blood at 0, 5, 15, or 30 min for insulin Insulin tolerance tests (ITT) were performed as described previously. Briefly, animals were fasted for 4h (09:00-13:00) and administered insulin (Novolin, 0.75 units per body weight: i.p.). Glucose was measured in tail blood by glucometer. Pyruvate tolerance tests (PTT) were performed as described previously. Briefly, animals were fasted for 6h (09:00-15:00) and administered pyruvate (2g/kg, i.p.; Sigma). Glucose was measured in tail blood by glucometer.

### Pancreatic islet isolation and in vitro assessment of insulin secretion

Islets were isolated by the Vanderbilt Islet and Pancreas Analysis (IPA) core by injecting 0.6 mg/ml collagenase P (Roche) into the pancreatic bile duct followed by a Histopaque-1077 (Sigma) fractionation and hand-picking. Isolated islets were cultured were cultured overnight in low glucose DMEM (Thermo, 11966-025) containing 1 g/L glucose and supplemented with 10% FBS (Atlanta Biologicals), 4.6 mM HEPES, 1% penicillin/streptomycin (Thermo), 1% non-essential amino acids (Sigma) in a sterile cell incubator at 37 °C with 5% CO2 infusion and 95% humidity. Glucose-stimulated insulin secretion (GSIS) was assessed by perifusion with the use of size-matched islets and normalized to islet equivalents as previously described(64). Briefly, three secretagogues, glucose (16.7 mmol/L), 3-isobutyl-1methylxanthine (IBMX) (100 μmol/L), and KCL (20mml/L), were used during perifusion. Insulin in the culture medium was determined by ELISA. Islet size was assessed with MetaMorph, version 7.7 (Universal Imaging).

### Immunohistochemistry and morphometric analysis

Anesthetized mice were perfused with saline and 4% paraformaldehyde in 100mM phosphate buffer, and their brains were dissected, postfixed in the same fixative overnight (4°C), and immersed in 30% sucrose. Hypothalami tissues were then separated from other regions, embedded in optimal cutting temperature compound, immediately frozen on dry ice, and stored at -80°C. Next, 35-μM-thick hypothalamic coronal sections were cut on a cryostat, blocked for 1 hour with 5% BSA in PBS containing 0.1% Triton X-100, and incubated with primary antibodies overnight at 4°C. Pancreata tissues were embedded in paraffin wax and sectioned to 6-μM thickness. Islet β- and α-cell areas were determined as described previously(65). Briefly, pancreatic sections taken every 50 μm (n=5 animals per condition) were scanned using a ScanScope CS scanner (Aperio Technologies, Vista, CA). Images from each experiment were processed with ImageScope Software (Aperio Technologies, Vista, CA). Islet b- and a-cell areas were calculated as the ratio of the insulin-or glucagon-positive area to the total pancreas area (eosin stained), Primary anti-mouse antibodies were against Iba1 (1:500, rabbit polyclonal, Wako, and 1:500, goat polyclonal, Abcam), CD68 (1:500, rat, Biorad) PSD95 (1:500, rabbit polyclonal, Synaptic Systems) Wisteria floribunda lectin (WFA, 1:500, Biotinylated, Vector Laboratories), Aggrecan (1:500, rabbit polyclonal, Millipore) AgRP (1:500, goat, Neuromics), GFP (1:500, chicken, Aves Labs), insulin (1:250, guinea pig, Fitzgerald industries), glucagon (1:100, rabbit, Cell Signaling) and somastotatin (1:500, goat, Santa Cruz Biotechnology). Adequate AlexaFluor-conjugated secondary antibodies were used for immunofluorescence microscopy. Sections were mounted with DAPI Vectashield solution (Vector Laboratories, Inc.) to identify cell nuclei. Images were acquired with a confocal laser-scanning microscope (Leica TCS SP8).

### Cell counting and analysis of microglial engulfment

Iba1+ cells in hypothalamic sections were counted manually by visual inspection of anatomically matched within prespecified regions of interest, with an average taken from three sequential sections per mouse. Microglial cell body size was determined by ImageJ software using a thresholding protocol (ImageJ) followed by densitometric quantification. WFA fluorescence intensity was similarly measured with Image J software from images captured using identical exposure times that also avoided saturating the pixel intensities. Imaris software 9.5.1 was used to create surface renderings of individual microglia cells labeled with Iba1, incomplete or poorly labeled cells were excluded from analyses. CD68, PSD95, and aggrecan content within the microglia surface were quantified. The volume of phagocytes and engulfed synapses was normalized to cell volume.

### Statistics

We analyzed all data using GraphPad Prism version 8.0 software (San Diego, CA). Data are presented as mean ± SEM. Two groups were compared using an unpaired two-tail Student’s t-test. One-way ANOVAs were used to analyze data sets with more than two groups. Two-way ANOVAs were used to analyze data sets with two independent variables, followed by Bonferroni post-hoc adjustment. P values of less than 0.05 were considered significant.

## Author Contributions

M.V. designed and conducted the experiments, analyzed data, and prepared figures. M.V. first drafted the manuscript. E.R.M., S.B.M., A.F., R.T.C., and L.L., assisted with key experiments. T.P.B. assisted with the PRV tracing studies. R.N.L, and A.W.X., provided key reagents, technical support and expertise. M.V. and S.K.K. conceived of the studies, supervised their design and completion, and wrote the final manuscript. M.V. and S.K.K. prepared the final figures.

## Acknowledgements

This work was supported by Larry L. Hillblom Foundation (Start-Up grant to M.V.), the NIDDK (R01 DK134782-01 to M.V., and R01 DK103175-02 to S.K.K.). Metabolic phenotyping was assisted by the Nutrition Obesity Research Center at UCSF(DK098722). Islet isolation and functional analysis were performed using the Islet and Pancreas Analysis (IPA) Core supported by the Vanderbilt Diabetes Research Center (NIH grant DK20593). The authors would like to thank Dr. Jens Bruning (Cologne, Germany) for helpful technical advice and insights.

---

**Figure S1. Maternal high-fat diet during lactation does not alter cortical microglial morphology and reactivity status in the offspring.** A) Representative confocal images of postnatal cortical sections, showing equivalent number and cell body size of Iba1+ microglia in the offspring of dams fed a standard low-fat diet (LFD) or high-fat diet (HFD) during the lactation period. B) Quantification of cell number in A. C) Quantification of cell body size in A (n= 5-group). Values are mean ± SEM, scale bar, 20 μm.

**Figure S2. Microglial depletion during the neonatal period (P6-16) does not affect glucose tolerance in adult offspring of LFD-fed dams.** A) Equivalent body weight and B) glucose tolerance test (GTT) at 8 weeks of age in microglial-depleted pups from P6-16 when dams were fed a low-fat diet during the lactation period. C) Area under the curve (AUC) of B (n=8-10/group). Values are mean ± SEM.

**Figure S3. Microglial depletion after the neonatal period (P21-31) in the offspring of HFD-fed dams.** A) Experimental design to assess the impact of microglial depletion from P21-31 on glucose tolerance in adult offspring of HFD-fed dams during lactation. B) Representative hypothalamic sections showing a nearly complete loss of Iba1 signal, indicating microglial depletion and repopulation after PLX5622 withdrawal (P36). Values are mean ± SEM (3V, third ventricle: scale bar, 20 μm).

**Figure S4. Mice treated with a CSF1R antagonist unable to cross the blood-brain barrier (PLX73086) during the neonatal period (P6-16) do not affect adult glucose tolerance in the offspring of HFD-fed dams.** A) Experimental design to assess the impact of PLX73086 from P6-16 on glucose tolerance in adult offspring of HFD-fed dams during lactation. B) Representative hypothalamic sections showing Iba1+ (green) microglial depletion with BLZ945 but not PLZ73086 at P16. C) Equivalent body weight and D) glucose tolerance test (GTT) in adult mice treated with PLX73086 from P6-16 when dams were fed a HFD during lactation. E) AUC of D (n=7-8/group). Values are mean ± SEM, scale bar, 20 μm.

**Figure S5. Microglial depletion during the neonatal period (P6-16) does not affect glucose tolerance in adult female offspring of HFD-fed dams.** A) Reduced body weight and B) equivalent glucose tolerance test (GTT) at 8 weeks of age in microglial-depleted female mice from P6-16 when dams were fed a high-fat diet during lactation. C) Area under the curve (AUC) of B (n=10-14/group). Values are mean ± SEM.

## References

1. Ruud J, Steculorum SM, Brüning JC. Neuronal control of peripheral insulin sensitivity and glucose metabolism. Nat Commun. 2017 May 4;8:15259.

2. Steculorum SM, Ruud J, Karakasilioti I, Backes H, Engström Ruud L, Timper K, et al. Agrp neurons control systemic insulin sensitivity via myostatin expression in brown adipose tissue. Cell. 2016 Mar 24;165(1):125–38.

3. Faber CL, Deem JD, Campos CA, Taborsky GJ, Morton GJ. CNS control of the endocrine pancreas. Diabetologia. 2020 Oct;63(10):2086–94.

4. Carey M, Kehlenbrink S, Hawkins M. Evidence for central regulation of glucose metabolism. J Biol Chem. 2013 Dec 6;288(49):34981–8.

5. Yoon NA, Diano S. Hypothalamic glucose-sensing mechanisms. Diabetologia. 2021 May;64(5):985–93.

6. Rosario W, Singh I, Wautlet A, Patterson C, Flak J, Becker TC, et al. The Brain-to-Pancreatic Islet Neuronal Map Reveals Differential Glucose Regulation From Distinct Hypothalamic Regions. Diabetes. 2016 Sep;65(9):2711–23.

7. Papazoglou I, Lee J-H, Cui Z, Li C, Fulgenzi G, Bahn YJ, et al. A distinct hypothalamus-to-β cell circuit modulates insulin secretion. Cell Metab. 2022 Feb 1;34(2):285–298.e7.

8. Li M, Sloboda DM, Vickers MH. Maternal obesity and developmental programming of metabolic disorders in offspring: evidence from animal models. Exp Diabetes Res. 2011 Sep 28;2011:592408.

9. Gaillard R, Felix JF, Duijts L, Jaddoe VWV. Childhood consequences of maternal obesity and excessive weight gain during pregnancy. Acta Obstet Gynecol Scand. 2014 Nov;93(11):1085–9.

10. Grove KL, Allen S, Grayson BE, Smith MS. Postnatal development of the hypothalamic neuropeptide Y system. Neuroscience. 2003;116(2):393–406.

11. Bouret SG, Draper SJ, Simerly RB. Formation of projection pathways from the arcuate nucleus of the hypothalamus to hypothalamic regions implicated in the neural control of feeding behavior in mice. J Neurosci. 2004 Mar 17;24(11):2797–805.

12. Nilsson I, Johansen JE, Schalling M, Hökfelt T, Fetissov SO. Maturation of the hypothalamic arcuate agouti-related protein system during postnatal development in the mouse. Brain Res Dev Brain Res. 2005 Mar 31;155(2):147–54.

13. Ahima RS, Prabakaran D, Flier JS. Postnatal leptin surge and regulation of circadian rhythm of leptin by feeding. Implications for energy homeostasis and neuroendocrine function. J Clin Invest. 1998 Mar 1;101(5):1020–7.

14. Bouret SG, Draper SJ, Simerly RB. Trophic action of leptin on hypothalamic neurons that regulate feeding. Science. 2004 Apr 2;304(5667):108–10.

15. Kamitakahara A, Bouyer K, Wang C-H, Simerly R. A critical period for the trophic actions of leptin on AgRP neurons in the arcuate nucleus of the hypothalamus. J Comp Neurol. 2018 Jan 1;526(1):133–45.

16. Chen H, Simar D, Morris MJ. Hypothalamic neuroendocrine circuitry is programmed by maternal obesity: interaction with postnatal nutritional environment. PLoS ONE. 2009 Jul 16;4(7):e6259.

17. Kirk SL, Samuelsson A-M, Argenton M, Dhonye H, Kalamatianos T, Poston L, et al. Maternal obesity induced by diet in rats permanently influences central processes regulating food intake in offspring. PLoS ONE. 2009 Jun 11;4(6):e5870.

18. Vogt MC, Paeger L, Hess S, Steculorum SM, Awazawa M, Hampel B, et al. Neonatal insulin action impairs hypothalamic neurocircuit formation in response to maternal high-fat feeding. Cell. 2014 Jan 30;156(3):495–509.

19. Douglass JD, Ness KM, Valdearcos M, Wyse-Jackson A, Dorfman MD, Frey JM, et al. Obesity-associated microglial inflammatory activation paradoxically improves glucose tolerance. Cell Metab. 2023 Sep 5;35(9):1613–1629.e8.

20. Abegg K, Hermann A, Boyle CN, Bouret SG, Lutz TA, Riediger T. Involvement of Amylin and Leptin in the Development of Projections from the Area Postrema to the Nucleus of the Solitary Tract. Front Endocrinol (Lausanne). 2017 Nov 28;8:324.

21. Liberini CG, Borner T, Boyle CN, Lutz TA. The satiating hormone amylin enhances neurogenesis in the area postrema of adult rats. Mol Metab. 2016 Oct;5(10):834–43.

22. Lutz TA, Coester B, Whiting L, Dunn-Meynell AA, Boyle CN, Bouret SG, et al. Amylin selectively signals onto POMC neurons in the arcuate nucleus of the hypothalamus. Diabetes. 2018 May;67(5):805–17.

23. Lutz TA, Le Foll C. Endogenous amylin contributes to birth of microglial cells in arcuate nucleus of hypothalamus and area postrema during fetal development. Am J Physiol Regul Integr Comp Physiol. 2019 Jun 1;316(6):R791–801.

24. Rosin JM, Vora SR, Kurrasch DM. Depletion of embryonic microglia using the CSF1R inhibitor PLX5622 has adverse sex-specific effects on mice, including accelerated weight gain, hyperactivity and anxiolytic-like behaviour. Brain Behav Immun. 2018 Oct;73:682– 97.

25. Thion MS, Ginhoux F, Garel S. Microglia and early brain development: An intimate journey. Science. 2018 Oct 12;362(6411):185–9.

26. Bohlen CJ, Friedman BA, Dejanovic B, Sheng M. Microglia in brain development, homeostasis, and neurodegeneration. Annu Rev Genet. 2019 Dec 3;53:263–88.

27. Neniskyte U, Gross CT. Errant gardeners: glial-cell-dependent synaptic pruning and neurodevelopmental disorders. Nat Rev Neurosci. 2017 Nov;18(11):658–70.

28. Wilton DK, Dissing-Olesen L, Stevens B. Neuron-Glia Signaling in Synapse Elimination. Annu Rev Neurosci. 2019 Jul 8;42:107–27.

29. Crapser JD, Arreola MA, Tsourmas KI, Green KN. Microglia as hackers of the matrix: sculpting synapses and the extracellular space. Cell Mol Immunol. 2021 Nov;18(11):2472–88.

30. Nguyen PT, Dorman LC, Pan S, Vainchtein ID, Han RT, Nakao-Inoue H, et al. Microglial remodeling of the extracellular matrix promotes synapse plasticity. Cell. 2020 Jul 23;182(2):388–403.e15.

31. Reichelt AC, Hare DJ, Bussey TJ, Saksida LM. Perineuronal nets: plasticity, protection, and therapeutic potential. Trends Neurosci. 2019 Jul;42(7):458–70.

32. Wen TH, Binder DK, Ethell IM, Razak KA. The perineuronal “safety” net? perineuronal net abnormalities in neurological disorders. Front Mol Neurosci. 2018 Aug 3;11:270.

33. Mirzadeh Z, Alonge KM, Cabrales E, Herranz-Pérez V, Scarlett JM, Brown JM, et al. Perineuronal Net Formation during the Critical Period for Neuronal Maturation in the Hypothalamic Arcuate Nucleus. Nat Metab. 2019 Feb;1(2):212–21.

34. Sun J, Wang X, Sun R, Xiao X, Wang Y, Peng Y, et al. Microglia shape AgRP neuron postnatal development via regulating perineuronal net plasticity. Mol Psychiatry. 2023 Nov 24;

35. Schafer DP, Lehrman EK, Kautzman AG, Koyama R, Mardinly AR, Yamasaki R, et al. Microglia sculpt postnatal neural circuits in an activity and complement-dependent manner. Neuron. 2012 May 24;74(4):691–705.

36. 36. Weinhard L, di Bartolomei G, Bolasco G, Machado P, Schieber NL, Neniskyte U, et al. Microglia remodel synapses by presynaptic trogocytosis and spine head filopodia induction. Nat Commun. 2018 Mar 26;9(1):1228.

37. Valdearcos M, Douglass JD, Robblee MM, Dorfman MD, Stifler DR, Bennett ML, et al. Microglial inflammatory signaling orchestrates the hypothalamic immune response to dietary excess and mediates obesity susceptibility. Cell Metab. 2017 Jul 5;26(1):185–197.e3.

38. Bellver-Landete V, Bretheau F, Mailhot B, Vallières N, Lessard M, Janelle M-E, et al. Microglia are an essential component of the neuroprotective scar that forms after spinal cord injury. Nat Commun. 2019 Jan 31;10(1):518.

39. 39. Cianfarani S, Germani D, Branca F. Low birthweight and adult insulin resistance: the “catch-up growth” hypothesis. Arch Dis Child Fetal Neonatal Ed. 1999 Jul;81(1):F71–3.

40. Veening MA, Van Weissenbruch MM, Delemarre-Van De Waal HA. Glucose tolerance, insulin sensitivity, and insulin secretion in children born small for gestational age. J Clin Endocrinol Metab. 2002 Oct;87(10):4657–61.

41. Cottrell EC, Ozanne SE. Early life programming of obesity and metabolic disease. Physiol Behav. 2008 Apr 22;94(1):17–28.

42. Bouret SG. Nutritional programming of hypothalamic development: critical periods and windows of opportunity. Int J Obes Suppl. 2012 Dec 11;2(Suppl 2):S19–24.

43. Thaler JP, Yi C-X, Schur EA, Guyenet SJ, Hwang BH, Dietrich MO, et al. Obesity is associated with hypothalamic injury in rodents and humans. J Clin Invest. 2012 Jan;122(1):153–62.

44. Valdearcos M, Robblee MM, Benjamin DI, Nomura DK, Xu AW, Koliwad SK. Microglia dictate the impact of saturated fat consumption on hypothalamic inflammation and neuronal function. Cell Rep. 2014 Dec 24;9(6):2124–38.

45. Weisberg SP, McCann D, Desai M, Rosenbaum M, Leibel RL, Ferrante AW. Obesity is associated with macrophage accumulation in adipose tissue. J Clin Invest. 2003 Dec;112(12):1796–808.

46. Liang W, Qi Y, Yi H, Mao C, Meng Q, Wang H, et al. The roles of adipose tissue macrophages in human disease. Front Immunol. 2022 Jun 9;13:908749.

47. Fujita Y, Nakanishi T, Ueno M, Itohara S, Yamashita T. Netrin-G1 Regulates Microglial Accumulation along Axons and Supports the Survival of Layer V Neurons in the Postnatal Mouse Brain. Cell Rep. 2020 Apr 28;31(4):107580.

48. Ozdinler PH, Macklis JD. IGF-I specifically enhances axon outgrowth of corticospinal motor neurons. Nat Neurosci. 2006 Nov;9(11):1371–81.

49. Sakai J. Core Concept: How synaptic pruning shapes neural wiring during development and, possibly, in disease. Proc Natl Acad Sci USA. 2020 Jul 14;117(28):16096–9.

50. Scott-Hewitt N, Perrucci F, Morini R, Erreni M, Mahoney M, Witkowska A, et al. Local externalization of phosphatidylserine mediates developmental synaptic pruning by microglia. EMBO J. 2020 Aug 17;39(16):e105380.

51. Lehrman EK, Wilton DK, Litvina EY, Welsh CA, Chang ST, Frouin A, et al. CD47 Protects Synapses from Excess Microglia-Mediated Pruning during Development. Neuron. 2018 Oct 10;100(1):120–134.e6.

52. Crapser JD, Spangenberg EE, Barahona RA, Arreola MA, Hohsfield LA, Green KN. Microglia facilitate loss of perineuronal nets in the Alzheimer’s disease brain. EBioMedicine. 2020 Aug;58:102919.

53. Crapser JD, Ochaba J, Soni N, Reidling JC, Thompson LM, Green KN. Microglial depletion prevents extracellular matrix changes and striatal volume reduction in a model of Huntington’s disease. Brain. 2020 Jan 1;143(1):266–88.

54. Liu Y-J, Spangenberg E, Tang B, Holmes TC, Green KN, Xu X. Microglia elimination increases neural circuit connectivity and activity in adult mouse cortex. J Neurosci. 2020 Dec 23;

55. Alonge KM, Mirzadeh Z, Scarlett JM, Logsdon AF, Brown JM, Cabrales E, et al. Hypothalamic perineuronal net assembly is required for sustained diabetes remission induced by fibroblast growth factor 1 in rats. Nat Metab. 2020 Oct;2(10):1025–33.

56. Chen W, Mehlkop O, Scharn A, Nolte H, Klemm P, Henschke S, et al. Nutrient-sensing AgRP neurons relay control of liver autophagy during energy deprivation. Cell Metab. 2023 May 2;35(5):786–806.e13.

57. Dearden L, Bouret SG, Ozanne SE. Sex and gender differences in developmental programming of metabolism. Mol Metab. 2018 Sep;15:8–19.

58. Dearden L, Balthasar N. Sexual dimorphism in offspring glucose-sensitive hypothalamic gene expression and physiological responses to maternal high-fat diet feeding. Endocrinology. 2014 Jun;155(6):2144–54.

59. Lenz KM, McCarthy MM. A starring role for microglia in brain sex differences. Neuroscientist. 2015 Jun;21(3):306–21.

60. Schwarz JM, Sholar PW, Bilbo SD. Sex differences in microglial colonization of the developing rat brain. J Neurochem. 2012 Mar;120(6):948–63.

61. Nelson LH, Warden S, Lenz KM. Sex differences in microglial phagocytosis in the neonatal hippocampus. Brain Behav Immun. 2017 Aug;64:11–22.

62. Huang Y, Xu Z, Xiong S, Sun F, Qin G, Hu G, et al. Repopulated microglia are solely derived from the proliferation of residual microglia after acute depletion. Nat Neurosci. 2018 Apr;21(4):530–40.

63. Milinkeviciute G, Henningfield CM, Muniak MA, Chokr SM, Green KN, Cramer KS. Microglia regulate pruning of specialized synapses in the auditory brainstem. Front Neural Circuits. 2019 Aug 28;13:55.

64. 64. and Pancreas Analysis Core I. Mouse Islet Perifusion (3-stimuli protocol) v1. 2022 Jan 12;

65. Cyphert HA, Walker EM, Hang Y, Dhawan S, Haliyur R, Bonatakis L, et al. Examining How the MAFB Transcription Factor Affects Islet β-Cell Function Postnatally. Diabetes. 2019 Feb;68(2):337–48.

