## Supplementary figures and images for "Microglia mediate the early-life programming of adult glucose control"

### Supplemental Figure 1

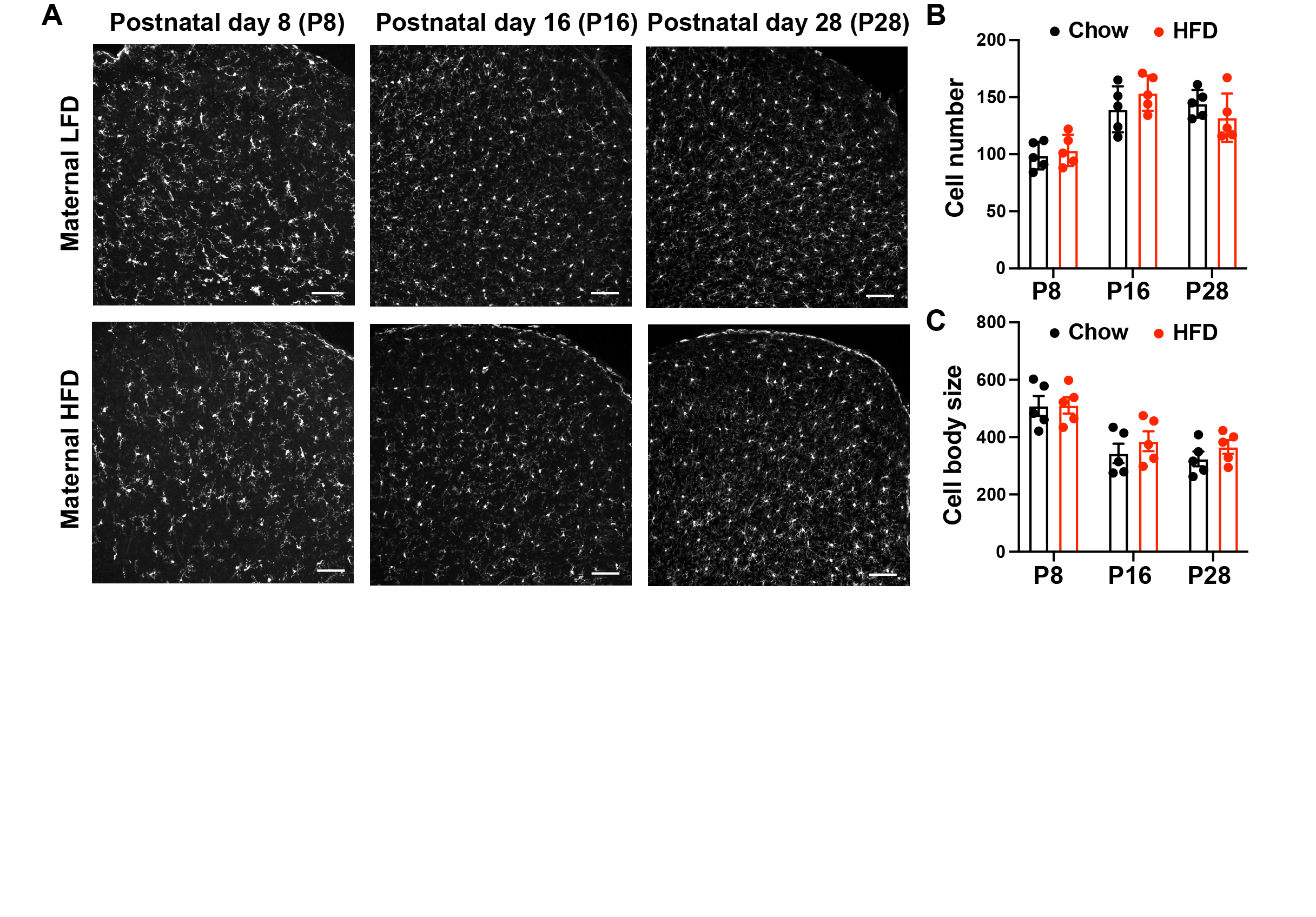

### Supplemental Figure 2

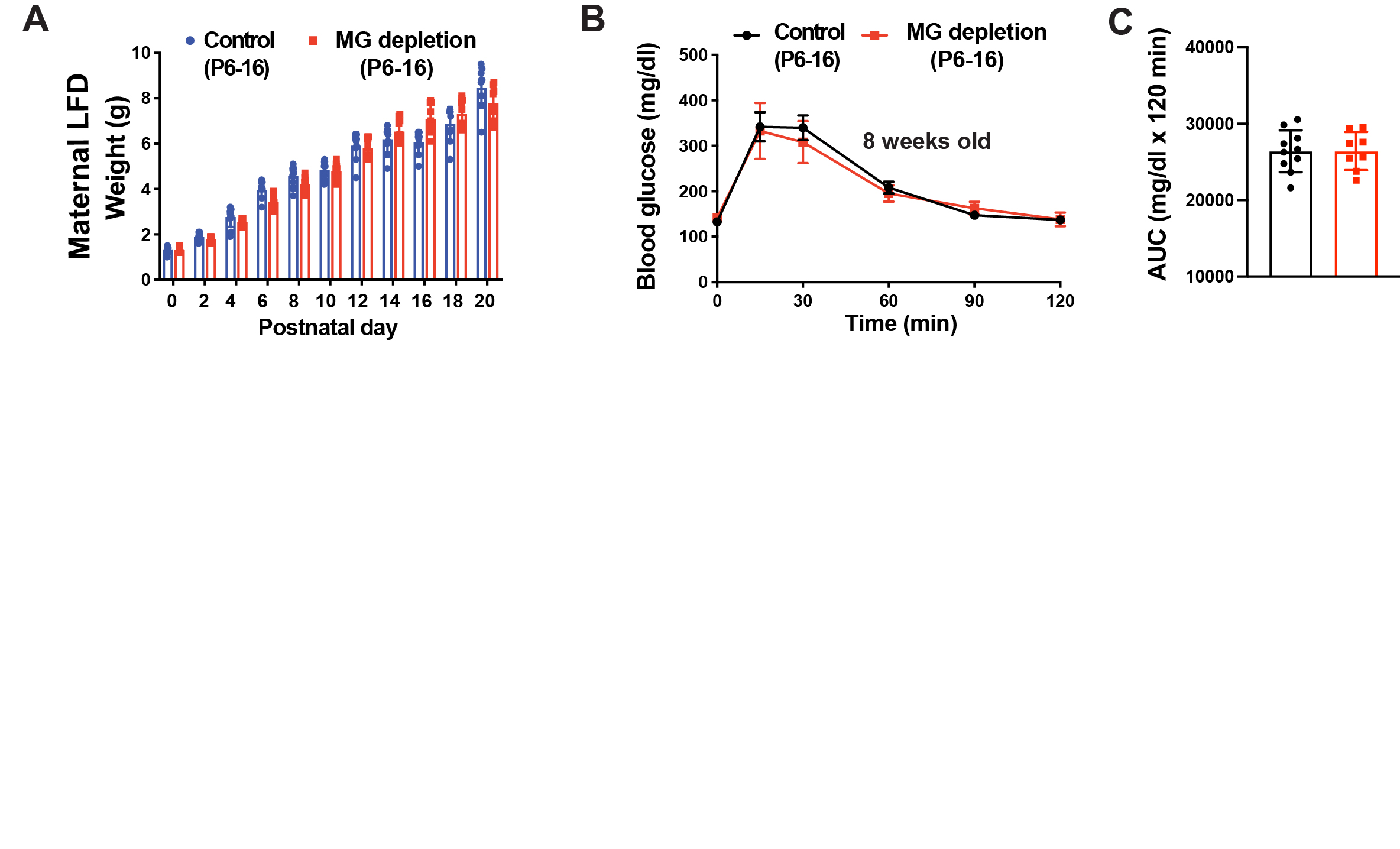

### Supplemental Figure 3

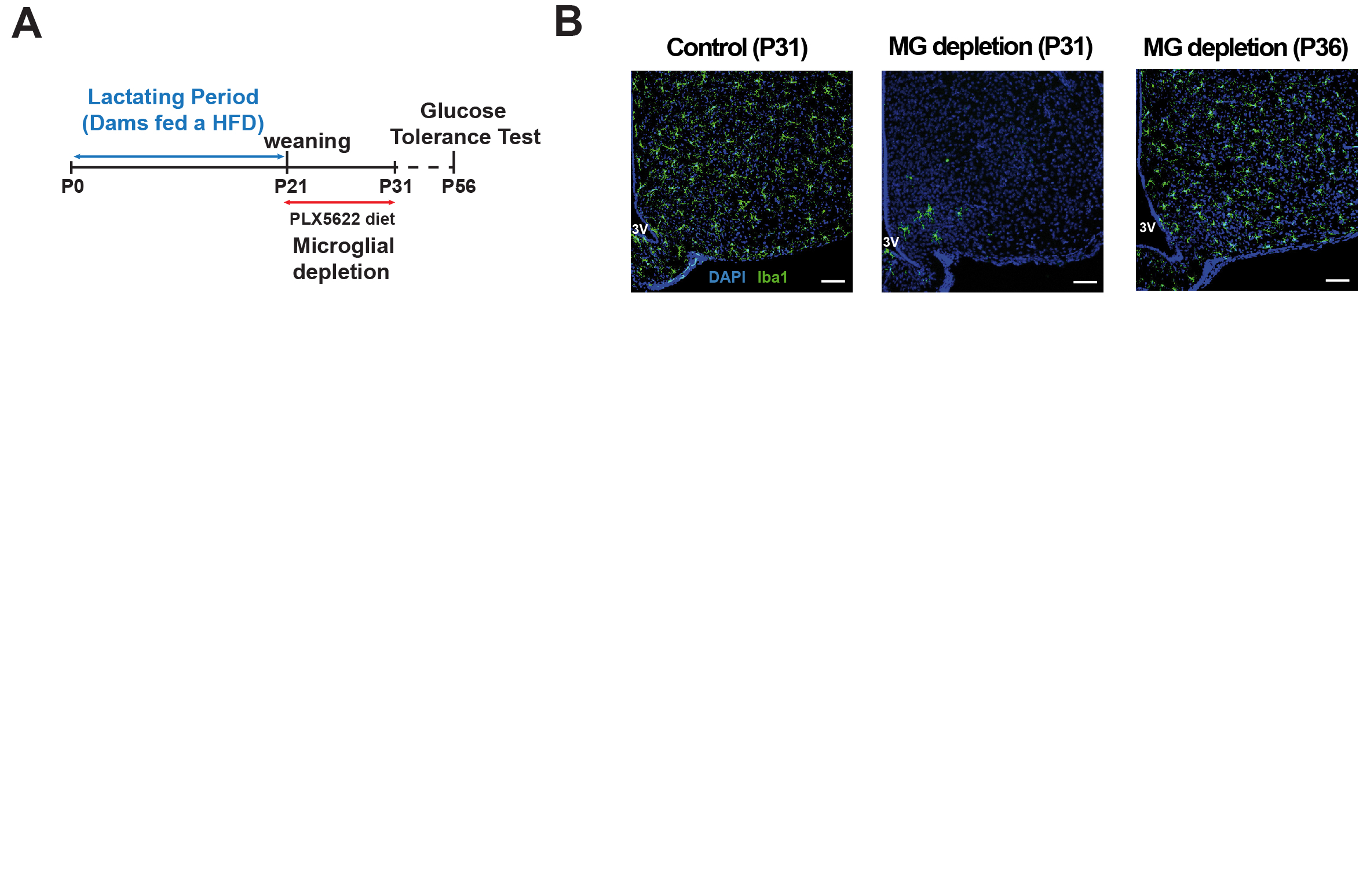

### Supplemental Figure 4

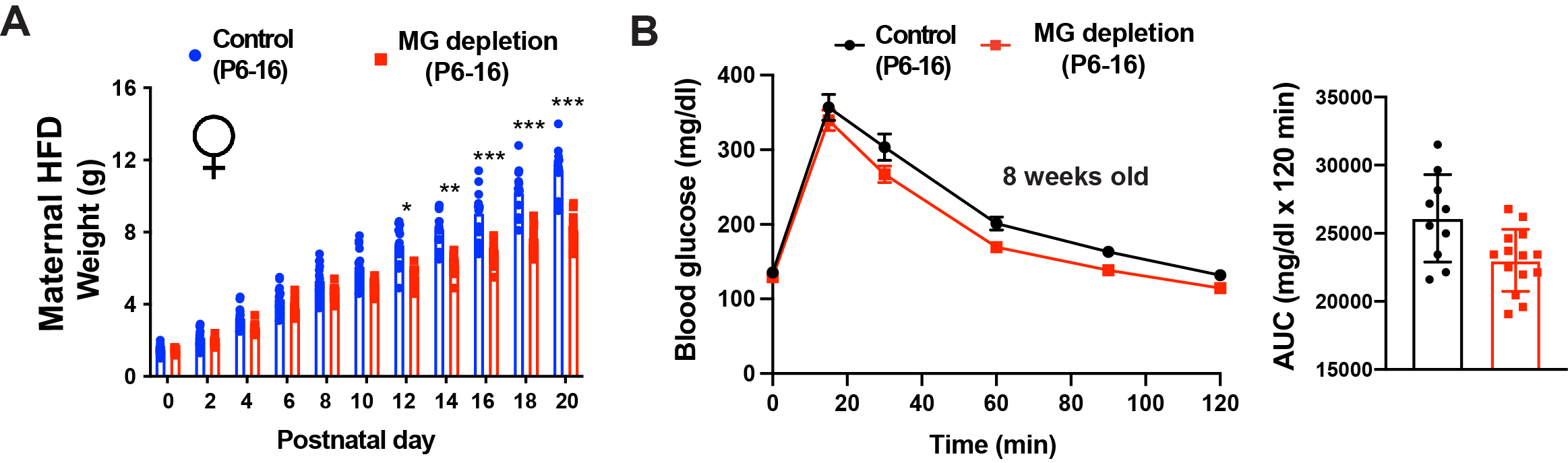
